# Atypical antipsychotics alter microglial functions via astrocyte-derived extracellular vesicles

**DOI:** 10.1101/2025.05.28.656664

**Authors:** Hana Yeh, Yoonjae Song, Liam T. McCrea, Joshua J. Bowen, Steven D. Sheridan, Roy H. Perlis

## Abstract

A limited understanding of the underlying molecular mechanisms of atypical antipsychotics has hindered efforts to develop the next generation of treatments for schizophrenia. In particular, there has been little investigation of how medications like clozapine and olanzapine modulate human non-neuronal cells, including astrocytes and microglia. Recent postmortem and serum-based studies suggest that schizophrenia etiology involves dysregulated cellular communication through extracellular vesicles (EVs). Astrocytes are a major source of these EVs and are strongly implicated in the etiology of schizophrenia by convergent data from human postmortem, brain imaging, RNA-sequencing, and genome-wide association studies. We hypothesized that clozapine and olanzapine can affect microglia biology indirectly via astrocytic secretion of EVs. We used *in vitro* cellular models with primary human astrocytes and PBMC-derived microglial-like cells to investigate the downstream impact of isolated astrocyte-derived EVs (ADEVs) on microglial phenotypes relevant to schizophrenia, including microglial phagocytosis, motility, and morphology. To model microglia-mediated synaptic pruning *in vitro*, we utilized image-based quantification of microglia engulfment of isolated human synaptosomes. We found that treatment with ADEVs reduced microglial synaptosome phagocytosis in a dose-dependent manner. This reduction was reversed upon addition of ADEVs isolated from astrocytes treated with norclozapine or olanzapine. ADEVs isolated from clozapine-treated astrocytes increased microglial motility, indicating that clozapine alters microglial surveillance activity without affecting phagocytosis through these ADEVs. Together, these results suggest that atypical antipsychotics have distinct and indirect impact on microglia biology mediated by ADEVs. These results highlight a potentially critical role for ADEVs in regulating glial cell communication and suggest they may be promising therapeutic targets for next-generation antipsychotic development.

## Introduction

Schizophrenia is a chronic and debilitating disorder with limited treatment options. Atypical antipsychotics, which target both dopamine and serotonin receptors in the brain, are not effective for all patients and carry substantial risk for adverse effects^1^. About 30-40% of individuals with schizophrenia do not respond adequately to multiple first-line antipsychotics, such as olanzapine, and become eligible for clozapine, the only FDA-approved antipsychotic for treatment-resistant schizophrenia patients^2^, and the only drug shown to diminish suicide risk in a randomized trial^3^. Though the efficacy of clozapine is greater than other atypical antipsychotics in those patients who do not respond to initial treatments, the molecular mechanisms and cellular specificity underlying its effectiveness remain unclear^2,4^. Further, the complexity of antipsychotics’ effects on non-neuronal cells complicates efforts to address these crucial gaps in mechanistic knowledge. In particular, the specific effects of clozapine and olanzapine on glial cells, such as astrocytes and microglia, remain relatively unknown, even though our previous work^5^ and others^6–8^ support the likelihood that non-neuronal cell types play a crucial role in schizophrenia pathophysiology.

Recent proteome, transcriptome and genome-wide studies support the notion that astrocytes contribute to schizophrenia pathogenesis via their role in multiple brain homeostasis mechanisms^9^. Astrocytes are important modulators of immune response, synaptic modulation and energy modulation^8^. Studies using iPSC-derived astrocytes demonstrated enhanced astrocytic calcium activity, reduced glucose uptake capacity, and dysregulated inflammatory profile associated with disrupted vascularization in schizophrenia patients-derived cells, further supporting a role for astrocytes in the etiology of schizophrenia^8,10,11^. Notably, schizophrenia patient-derived astrocyte cell transplantation to the mouse forebrain caused gene expression changes in synaptic dysfunction and inflammation pathways of mouse cortical cells, resulting in behavioral changes in cognitive and olfactory functions^6^.

A particular gap in knowledge is the impact of atypical antipsychotics on astrocytes or microglia, much less on the cellular communication between these glial cells. One study found clozapine rescued deficient levels of glutamate and NMDA receptor coagonist D-serine levels in astrocytes derived from clozapine-responsive schizophrenia patients^12^. Another study found that clozapine metabolites including norclozapine inhibited microglial NADPH oxidase to protect dopaminergic neurons^13^. How antipsychotic drugs alter the interaction between microglia and astrocytes is unknown.

One means by which neural cells interact relies on extracellular vesicles (EVs), secreted by neurons and glial cells, including astrocytes and microglia, with demonstrated roles in neuron-neuron, glia-neuron, and glia-glia communication^14,15^. EVs are lipid membrane-contained vesicles carrying proteins, lipids, and nucleic acids, secreted by cells for intercellular communication^16^. Recent studies have shown altered protein expression in astrocyte-derived EVs (ADEVs) from individuals with schizophrenia, including increased inflammatory-associated proteins and dysregulated expression of mitochondrial proteins following the first psychotic episode, suggesting astroglia pathology^17–19^. ADEVs secreted from non-stimulated astrocytes show neurotrophic and neuroprotective properties, whereas ADEVs released from astrocytes in ischemic, oxidative stress, nutrient-deprived, or thermal stress conditions are enriched with proteins and microRNAs (miRNAs) that promote neuronal survival, reduce neurotransmitter toxicity, and enhance neurite outgrowth^20–25^. Recent studies show that EVs can also contribute to the pathology of psychiatric disorders through modulation of miRNA content^26,27^. Serum-derived EVs from schizophrenia patients injected into mice result in schizophrenia-like behavior and transcriptomic changes related to neurodevelopment dysregulation^28^. Therefore, investigating the role of ADEVs can show insight into glial cellular communication.

Thus, despite multiple lines of evidence suggesting the importance of astrocytes as well as EVs in psychotic disorders, the possibility that antipsychotics modulate microglial function via ADEVs remains to be explored. We hypothesized that ADEVs could regulate microglia and that these effects could be moderated by antipsychotic exposure. In light of postmortem and clinical structural neuroimaging studies demonstrating reduced cortical dendritic spine density among individuals with schizophrenia, a particular disease target may be microglia-mediated synaptic pruning dysfunction^5,29–31^. Here, we show that different atypical antipsychotics have distinct impact on various microglial biology including phagocytosis, motility and morphology, and that such impacts can be mediated by ADEVs.

## Methods

### Ethical Statement

The human samples investigated in this study were collected under a tissue banking protocol #2019P001196 approved by the Mass General Brigham Institutional Review Board (IRB), with informed consent obtained from all participants. The present study used de-identified samples from an IRB-approved tissue bank.

### Primary human astrocytes and drug treatment

Human primary astrocytes (HA1800) were purchased from ScienCell and tested negative for mycoplasma. HA1800 cells were plated on poly-L-lysine-coated T-75 and T-175 flasks and passaged with TrypLE Express (Gibco, # 12605010). HA1800 cells were cultured in complete Astrocyte Media (ScienCell, #1801). Media change was performed every 2-4 days and passaged using TrypLE Express for passage 4-5 experimental assays. For immunocytochemistry, HA1800 cells were plated onto 96-well optical plates (Corning, #3904) at 10,000 cells per well density. Primary astrocytes were passaged and plated onto 96-well (ThermoScientific, #165306) at 10,000 per well density or 10 million cells per T-75 flask for EV harvest. The antipsychotic drugs clozapine (Sigma, #C6305), norclozapine (Sigma, #D5676) and olanzapine (Sigma, #O1141) were dissolved in DMSO (Sigma, #D2438). Serial dilution in complete Astrocyte Media was performed to final concentration of 300 μM, 150 μM, 75 μM, 37.5 μM, 18.8 μM, and 9.4 μM. Following 48 hrs, CellTiter-Glo Luminescent Cell Viability Assay (Promega, #G7571) was performed to determine optimal concentration with EnVision Multilabel reader.

### Astrocyte-derived extracellular vesicle (ADEV) generation and isolation

Human astrocytes were passaged and plated into T-75 flasks at 10 million per flask in complete Astrocyte Media (ScienCell, #1801). After two days, media was switched to exosome-depleted astrocyte basal media consisting of DMEM/F12 (Gibco, #11320033), 2% exosome-depleted (Gibco, #A2720803), 1X N2, 1X GlutaMax, 1X Pen/Strep, and 1X MEM non-essential amino acids. 20 ng/ml IL1β (Peprotech, #200-01B) and 500 μM adenosine triphosphate (ATP, Sigma, #A7699-1G) were directly added T-75 flasks. Clozapine, norclozapine, and olanzapine were added to a final concentration of 30 µM. Media were collected 48 hours post-treatment. ADEV-containing media was centrifuged at 300 g for 10 minutes at 4°C to remove dead cells, then at 4,000 g for 30 minutes at 4°C to remove debris. The media was filtered using a 0.22 µm PES SteriFlip Filter (EMD Millipore, #SCGP00525). ADEVs were isolated using Total Exosome Isolation Reagent (Invitrogen, #4478359) and incubated overnight at 4°C. Precipitated ADEVs were recovered by centrifugation at 10,000 g for 60 minutes and resuspended in double-filtered PBS or exosome-resuspension buffer (Invitrogen, #4478545). The isolated ADEVs were further washed and centrifuged at 100,000 g for 2 hours at 4°C for RNA extraction kit (Invitrogen, #4478545) or aliquoted for storage at −80°C ^32^.

### ADEV Nanoparticle Tracking Analysis

For nanoparticle tracking analysis to determine EV concentration and size distribution, ADEVs were thawed on ice and resuspended in double-filtered PBS^33^. ADEVs were analyzed using Nanosight LM10 (Malvern) equipped with an AVT MARLIN F-033B IRF camera (Allied Vision Technologies) and NTA 3.1 Build 3.1.46 software. All nanoparticle tracking analyses were carried out with identical experiment settings (Camera Level = 11, Detection Threshold = 7). Particles were measured for 30 s and each sample was imaged with at least 4 technical replicates. Videos were analyzed using NTA 3.2 software (Malvern).

### ADEV Protein Quantification and Western blot

For protein extraction, ADEVs were lysed with RIPA buffer containing a protease inhibitor cocktail (Millipore Sigma, #11836170001), incubated on ice for 10 minutes, and centrifuged at 15,000 RPM for 10 minutes at 4°C. The supernatant was collected, and protein concentration was determined using the Pierce BCA Protein Assay (Thermo Scientific, #23225). Protein samples (5 µg) were mixed with 1x Loading Dye (Boston BioProducts, #BP111R) and heated to 95°C for 2 minutes. For CD9 and CD81 antibodies, β-mercaptoethanol (B-ME) was omitted, while 355 mM B-ME was added to other samples. Precision Plus Protein Dual Color Standards (Bio-Rad, #1610374) were used for ladders. Samples were run on a mini-PROTEAN gel (Bio-Rad, #456-1096) in 1x Tris-Glycine-SDS running buffer (Boston BioProducts, #BP-150) at 60 V for 15 minutes, followed by 100 V for 1.5 hours. Proteins were transferred to an Immun-Blot PVDF membrane (Bio-Rad, #1620177) using the BioRad Mini PROTEAN Tetra Cell system. Transfer was conducted at 125 V for 1 hour. Membranes were blocked with LICOR Intercept Blocking Buffer (LI-COR, #927-70001) for 1 hour at room temperature. Primary antibodies (Rabbit anti-CD9, Abcam, #ab263019; Rabbit anti-CD81, Abcam, #ab109201; Rabbit anti-TSG101, Abcam, #ab125011; Rabbit anti-Calnexin, Abcam, #ab133615) were diluted 1:1000 in Blocking Buffer and 0.2% Tween-20 (Sigma, #P2287) and incubated overnight at 4°C. Membranes were washed with PBS and 0.01% Tween-20. Secondary antibodies (IRDye 800CW Donkey anti-Rabbit IgG, LI-COR, #926-32213) were diluted 1:15,000 in 50% Blocking Buffer, 50% DPBS, 0.01% Tween-20, 0.02% SDS and incubated for 45 minutes at room temperature. Final washes were performed as described, and membranes were imaged using the LICOR Odyssey Clx.

### Differentiation of PBMC-derived microglia-like cells (piMGLCs)

Human PBMCs were isolated from a fresh leukopak (Hemacare) using a gradient protocol. The leukopak was layered on 1.077 g/mL density gradient media (DGM) and centrifuged at 400 g for 30 minutes at room temperature (brake off). PBMCs at the interface were collected and washed by centrifugation at 400 g for 10 minutes. Cells were cryopreserved in 10% DMSO/90% HI-FBS. For reprogramming to induced microglia-like cells (piMGLCs), samples were thawed in at 37°C and resuspended in RPMI-1640 (Sigma, #R8758) with 10% heat-inactivated fetal bovine serum (Sigma, #12306C), 1% Penicillin/Streptomycin (Life Technologies, # 15140-122), and 1% GlutaMax (Life Technologies, # 35050-061). Cells were washed by centrifugation at 300 g for 5 minutes and seeded at approximately 30K cells/cm² in Geltrex-coated (Gibco, #A1413202) plates. After 24 hours, the media was replaced with RPMI-1640 containing 1% Penicillin/Streptomycin, 1% GlutaMax, 100 ng/mL IL-34 (PeproTech, #200-34), and 10 ng/mL GMCSF (PeproTech, #300-03). Half of the media was changed at DIV6. PBMC-iMGLs were replated using Accutase at DIV8 to 15,000 cells per 96-well plate with 50% fresh media and differentiated until DIV14 for assays.

### Real-time live-imaging of microglia-like cell for motility tracking

Live imaging of microglia-like cellular cultures was performed using IncuCyte S5 (Sartorius) as previously described^34^. piMGLCs were imaged at 5-minute intervals for 5 hrs. total. Image sets were exported from the software as 8-bit .tif files and imported into FIJI image analysis software version 2.3.0/1.53q as image sequences. Image sequences were stabilized using the stabilization plug-in Image Stabilizer in ImageJ^35^. To track cells in the sequence, phase contrast image stacks were evaluated after ‘find edges’ processing on ImageJ with further processing using Trackmate 7 plug-in to trace and quantify individual cellular motility tracks (e.g. X, Y positions over time for each cell)^36^. Velocity (μm/min) and accumulated distance (μm from start of tracing) data were calculated using 0.61μm/pixel as a conversion factor for each track and additionally represented by motility plots using ImageJ Chemotaxis and Migration Tool plug-in^37^.

### Microglial synaptosome phagocytosis assay

Phagocytosis assays were conducted in 96-well plates (Corning, #3904) with microglia at a density of 12,000-15,000 cells per well as previously described^34^. Human synaptosomes were thawed at room temperature and mixed with an equal volume of 0.1M sodium bicarbonate (pH 9.0). pHrodo-Red (6.67 µg/µL, Invitrogen, #P36600) was added at a 1:2 dye-to-protein ratio and incubated for 1 hour in the dark. Synaptosomes were washed with PBS and centrifuged at 12,000 rpm for 15 minutes. The pellet was resuspended to a final concentration of 0.6 µg/µL and sonicated in a Branson 1800 (Emerson, #M1800) at 40 kHz for 1 hour. For positive control, microglia were treated with 10 μM cytochalasin-D (Sigma, #C2618) 1 hour before adding synaptosomes. Synaptosomes were added to microglia-like cells at 3 µg/well and incubated for 3 hours. The assay was terminated by fixing with 4% paraformaldehyde (Electron Microscopy Sciences, #15713S) for 15 minutes. The phagocytic index, defined as the area of pHrodo-Red signal divided by the area of IBA1+ cells per image, was calculated using RStudio to analyze metadata from CellProfiler (Version 4.2.1)^38^.

### Immunocytochemistry and confocal microscopy imaging

Live cells were fixed with 4% paraformaldehyde for 15 minutes at room temperature and washed three times with PBS. Cells in a 96-well format were then incubated with Block and Permeabilization Buffer (PBS, 0.5% FBS, and 0.3% Triton-X) for 1 hour at room temperature. Primary antibodies (Suppl. Table 1) were diluted in Antibody Buffer (PBS, 0.5% FBS, and 0.1% Triton-X), and incubated overnight at 4°C. After three washes with Wash Buffer (PBS and 0.5% FBS), secondary antibodies (Suppl. Table S1) and Hoechst 33352 (1:5000, ThermoFisher #62249) were diluted in Antibody Buffer were added and incubated for 45 minutes at 4°C. PKH67 dye (Sigma Aldrich, #P7333) was added according to vendor instructions. Following a final three washes, wells were imaged using an INCell Analyzer 6000 automated confocal microscope (Cytiva) at 20X magnification.

### Statistical analysis

Morphological, area, and intensity metrics were exported from CellProfiler for further analysis in RStudio. Graphing and statistical analyses were performed with R studio or GraphPad Prism 9 (GraphPad Software, La Jolla, CA USA, www.graphpad.com/scientific-software/prism). All reported *p*-values and type of statistical tests are reported in the figure legends.

## Results

### Reduction of PBMC-derived microglial phagocytosis, but not motility, by astrocyte-derived extracellular vesicles is dose-dependent

Previous studies have examined neuronal communication with glial cells such as microglia or astrocytes^39,40^, but not glial-to-glial communication via EVs, especially in psychiatric disorders. We investigated how astrocytes alter microglial biology via secreted EVs using human primary astrocytes and PBMC-derived microglia-like cellular models. In a dose-dependent manner, undiluted ADEVs (6.20E3±3.86E2 particles/cell ratio) reduce microglia phagocytosis, whereas diluted 1:10 ADEVs (6.20E2±3.86E1 particles/cell ratio) or further diluted concentrations had minimal impact on microglial phagocytosis, suggesting ADEV particles need to reach a critical threshold to alter microglial functions (Fig. 1A-B). Furthermore, microglial marker IBA1 expression following phagocytosis decreased significantly upon both undiluted and diluted 1:10 ADEV treatments, suggesting microglial biological responses are associated with inflammation in a dose-dependent manner (Fig. 1C), as relative IBA1 expression has been associated with inflammation or activation state^41^. Both reduced phagocytosis and expression of IBA1 suggest that ADEVs added at effective doses can dampen the inflammatory state and sensitivity to phagocytose synaptosomes.

**Figure 1.**
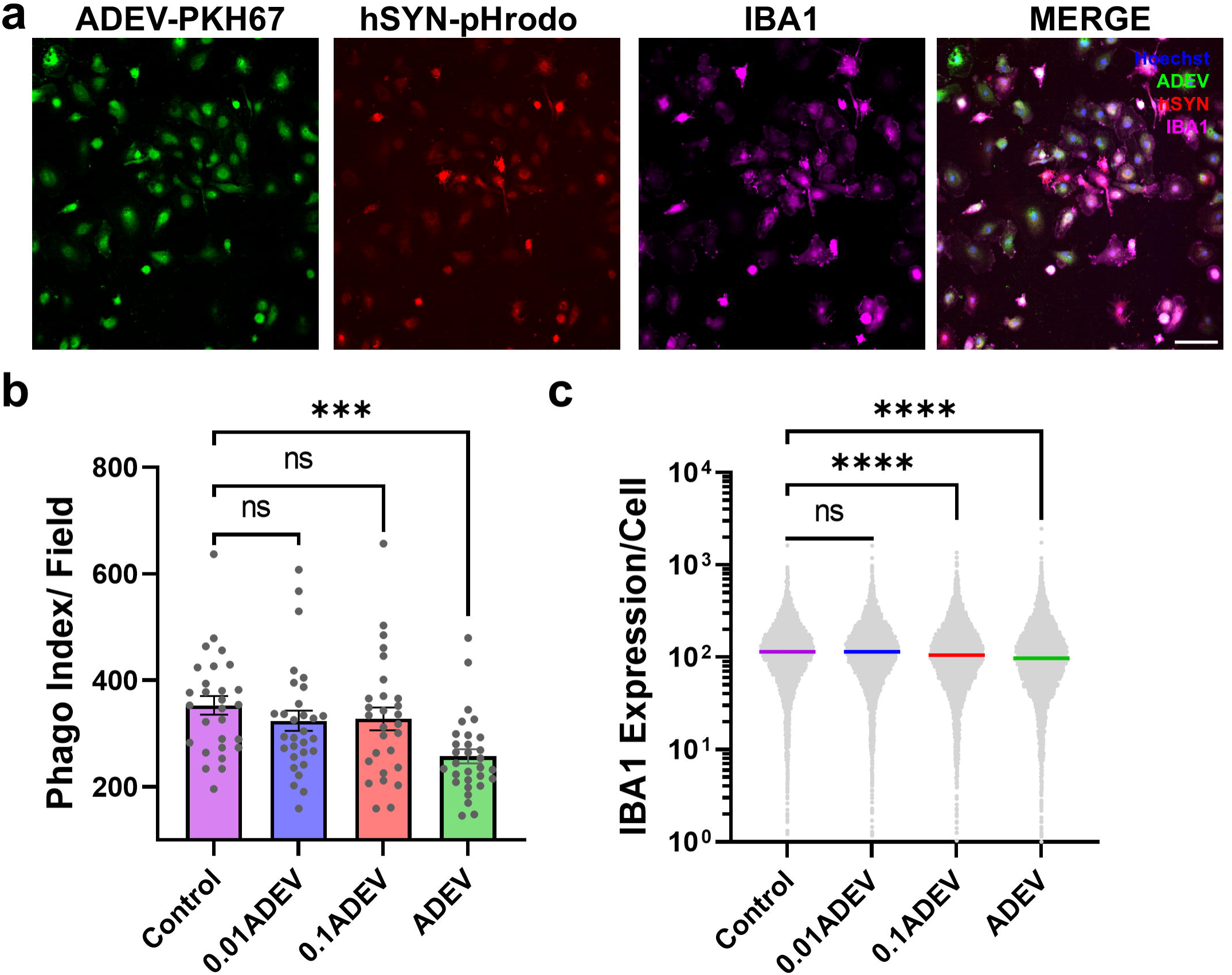
Dose-dependent reduction of microglial phagocytosis and IBA1 expression by ADEV treatment. **a**) Representative images for ADEVs in piMGLCs. Scale bar=50um. **b**) Phagocytosis index of piMGLCs. Each circle represents phagocytic index per image field. n=28-30 fields across 3 replicates. **c**) Reduced microglial marker IBA1 in a dose-dependent manner of ADEVs. n= 8448, 8192, 8900, and 8798 cells from 3 wells. Kruskal-Wallis test with Dunn’s multiple comparison test. Bars represent mean ± SEM, ∗∗∗p< .001, ∗∗∗∗p<.0001.

We next examined microglial motility, as they are the most motile cell type in the brain and such motility facilitates their capacity to shape circuitry during neurodevelopment and regulate immune response throughout adulthood. Microglia can also modulate synaptic activity through microglial contacts with dendritic spines^42–44^. Furthermore, ADEVs are enriched with proteins associated with regulation of actin cytoskeleton, which suggests they can alter the recipient cell cytoskeleton^45^. Here, we show that ADEV exposure does not alter microglial motility, suggesting microglial surveillance is independent of astrocytic-derived EV signaling (Suppl. Fig. 1).

### ADEVs isolated from ATP-stimulated astrocytes mediate microglia morphology and phagocytosis

Previous studies have shown that stimulation of glial cells alters secretion of EVs and their cargo content^46,47^. Astrocytes and microglia express purinergic receptors and readily respond to ATP, which is essential in directly activating astrocytes and eliciting microglia migration response in the brain parenchyma^47^. ATP can affect microglial physiology indirectly through EVs released from stimulated astrocytes. We treated astrocytes with ATP and harvested ADEVs 24 hours following treatment (ATP-ADEVs). We found ATP stimulation led to secretion of a larger proportion of ADEVs ranging from 200-400nm by nanoparticle tracking analysis (NTA) as indicated by increased ADEV mode size due to ATP-stimulation (Suppl. Fig. 2a-b). No significant EV concentration or size changes were observed due to ATP or IL1β stimulation (Suppl. Fig. 2b). We found that ATP-ADEV treatment led to increased ADEV load per cell based on area of PHK67-labeled ADEVs in IBA1+ microglia-like cells and increased cell area (Suppl. Fig. 2c-d).

Among the first indications of altered physiology in microglia are changes in microglia morphology, or cytoplasmic IBA1 and lysosomal CD68 expression. We further evaluated two critical morphologic variables, cell eccentricity and cell solidity, which can suggest the extent of ramified (low solidity, high eccentricity), bipolar (mid solidity, mid eccentricity) and amoeboid (high solidity, low eccentricity) morphology based on IBA1 segmentation using confocal immunofluorescence and CellProfiler quantification. When treated with ATP-ADEVs, microglial morphology was significantly altered, suggesting both functional and morphological changes that resemble an altered state including reduced IBA1 expression, more ramified, and larger cellular phenotype compared to CTRL group without ADEVs (Fig. 2a-c, Suppl. Fig. 2d).

**Figure 2.**
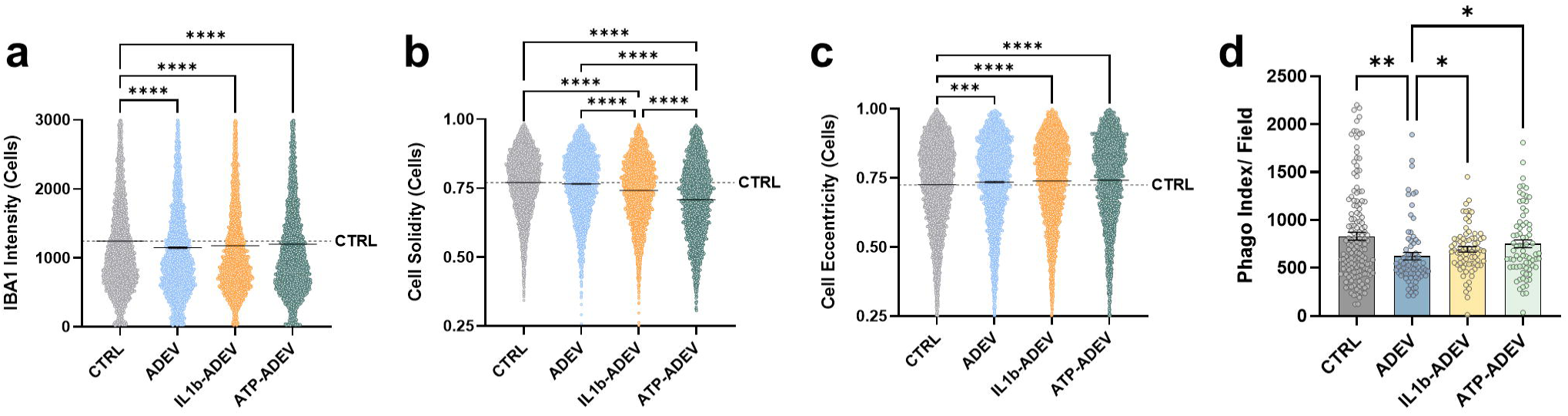
ADEV treatment alters microglial IBA1 expression, morphology, and phagocytosis. **a**) ADEV treatments reduced IBA1 expression in piMGLCs. **b**) IL1β-ADEVs and ATP-ADEVs reduced cell solidity on microglia morphology. **c)** ADEV treatment increased cell eccentricity morphology in piMGLCs. a-c) Each dot represented individual cell. Kruskal-Wallis test with Dunn’s multiple test (CTRL=4300, ADEV=4148, IL1β-ADEV=4300, ATP-ADEV=4390 cells). **d)** Synaptosome phagocytosis index reduced by ADEVs. Each dot represents per field. Kruskal-Wallis test with Dunn’s multiple test (CTRL=148 fields, ADEV, IL1β-ADEV, ATP-ADEV=75 fields). Bars represent mean ± SEM, ∗*p*<.05, ∗∗*p*<.01, ∗∗∗*p*<.001, ∗∗∗∗*p*<.0001.

We further investigated another stimulant to compare the impact of EVs isolated from astrocytes at different activation states. IL1β stimulation is known to induce inflammation, activate astrocytes, and increase neuronal excitability^48^. Proinflammatory IL1β cytokine treatment of astrocytes had minimal impact via ADEVs on microglial morphology. IBA1 expression and cell solidity were reduced by ADEVs, regardless of ATP or IL1β treatments (Fig. 2A-B). ADEV load per individual cell was determined based on PHK67-labeled ADEVs and divided into high versus low load to assess whether more ADEVs uptake per cell can alter microglial phenotype on the individual cell level in different treatment groups. In ADEV-treated groups, higher ADEV load per cell correlated with higher expression of IBA1 regardless of specific stimulation (Suppl. Fig. 2C, E). Both IL1β and ATP-treated ADEVs significantly reduced microglial cell solidity but had less impact on cell eccentricity (Fig. 2B-C) compared to ADEVs from untreated astrocytes– thus, ADEVs from both treatment conditions increased microglial ramification with concomitant reduced amoeboid morphology. While ADEVs reduced microglia phagocytosis under baseline conditions, ADEVs harvested after IL1β and ATP treatment had more modest effects on phagocytosis compared to control ADEVs (Fig. 2d). Together, these results suggest that altering ADEV cargo by different astrocyte stimulation in turn differentially affects microglia morphology and phagocytosis upon ADEV exposure.

### Atypical antipsychotics clozapine and olanzapine do not alter astrocytic marker expression or extracellular vesicle release from astrocytes

To determine viable dosage for clozapine and olanzapine treatment on human primary astrocytes *in vitro*, we used CellTitre Glo viability assay from 10μM to 300μM for 24 hours and compared to vehicle control (0.02% DMSO). Clozapine reduced cell viability when the concentration exceeded 32μM (Suppl. Fig. 3A). Norclozapine, also known as N-desmethylclozapine, is an active metabolite form of clozapine with distinct pharmacological properties^1,49^. Both clozapine and norclozapine were separately included as previous studies indicate they can have varying effects on cell biology, and norclozapine has been shown to increase dopamine and acetylcholine levels via muscarinic acetylcholine receptor M1 agonism, whereas clozapine act as M1 antagonist^50,51^. Based on immunofluorescence imaging, we confirmed that the antipsychotic treatments clozapine, norclozapine and olanzapine retained the expression of typical astrocyte markers, including GFAP, ALDH1L1, and S100b at 30μM concentration (Suppl. Fig. 3b). Thus, clozapine (CLOZ), norclozapine (NORC) and olanzapine (OLAN) were added to human primary astrocytes at 30μM for ADEV isolation (CLOZ-ADEV, NORC-ADEV and OLAN-ADEV, respectively), and compared to vehicle control (DMSO-ADEV).

Primary human astrocytes were treated 24 hours with atypical antipsychotics or vehicle control and the cell culture media were collected for extracellular vesicles. To characterize and validate antipsychotic-treated ADEVs, we performed western blot for extracellular markers including CD9, CD81, and TSG101, and non-extracellular marker calnexin to confirm quality of ADEVs isolation from primary astrocytic cell culture (Fig. 3A). We confirmed ADEV expression of both CD9 and CD81 regardless of antipsychotic treatment. The ADEV samples weakly expressed TSG101 (Fig. 3A). We further confirmed the size and concentration of ADEVs using NTA (Fig. 3B-C). CLOZ, NORC, and OLAN did not alter the extracellular vesicle concentration or mean size of extracellular vesicles. The mode size of the ADEVs suggests that a significant proportion of the EV sample resembles exosome size and contains larger size extracellular vesicles (Fig. 3C).

**Figure 3.**
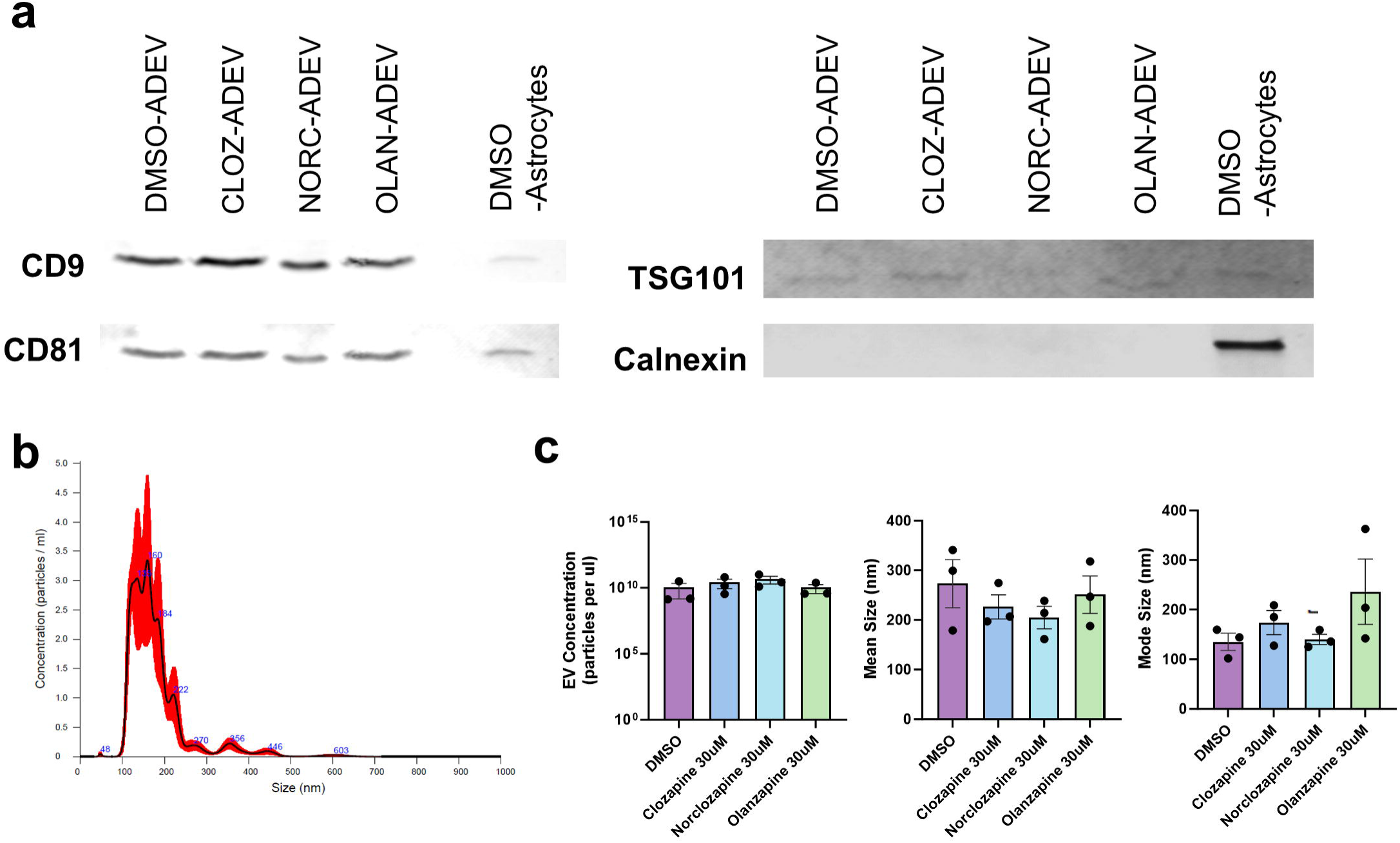
Atypical antipsychotics clozapine and olanzapine do not alter extracellular vesicle release from astrocytes. **a**) Atypical antipsychotics do not alter expression of extracellular vesicle markers CD9, CD81, and TSG101. All ADEV samples lack expression of cytoplasm marker calnexin by Western Blot. **b)** Representative ADEV trace from nanoparticle tracking analysis using Nanosight. **c)** No differences in mean concentration, mean size, and mode size observed by antipsychotic treatments. Mann-Whitney two-tailed test compared to DMSO treatment, *n*=3 replicates.

### The atypical antipsychotics norclozapine and olanzapine regulate microglial phagocytosis via ADEVs

Previous studies have shown that dysregulation of microglial phagocytosis can contribute to the pathophysiology of schizophrenia^5,52^, and our prior work demonstrated that microglia-like cells derived from individuals with schizophrenia show increased phagocytosis^5,34^. To study the impact of ADEV isolated from antipsychotic-treated astrocytes on microglial phagocytosis, we utilized our *in vitro* cellular model for human synaptosome phagocytosis with PBMC-derived microglia-like cells (piMGLCs)^5,53^ and measured differential effects on microglial phagocytosis upon exposure of ADEVs derived from antipsychotic treated astrocytes. Previous studies have shown that CLOZ can normalize microglia phagocytosis in presence of iron^54^. Here we show that ADEV isolated from OLAN- and NORC-treated astrocytes increased microglial synaptosome phagocytosis. (Fig. 4A-B). The increased phagocytosis index and phagocytosis index intensity show piMGLCs exposure to ADEVs derived from drug-treated astrocytes engulf more pHrodo-labelled synaptosomes compared to control DMSO-ADEV treated piMGLCs (Fig. 4A-B). Increased phagocytosis due to exposure to ADEVs derived from antipsychotic-treated astrocytes suggests antipsychotics can impact microglia functions via EVs secreted from astrocytes in an indirect manner, possibly due to different EV cargo that have different downstream effects on microglial biology.

**Figure 4.**
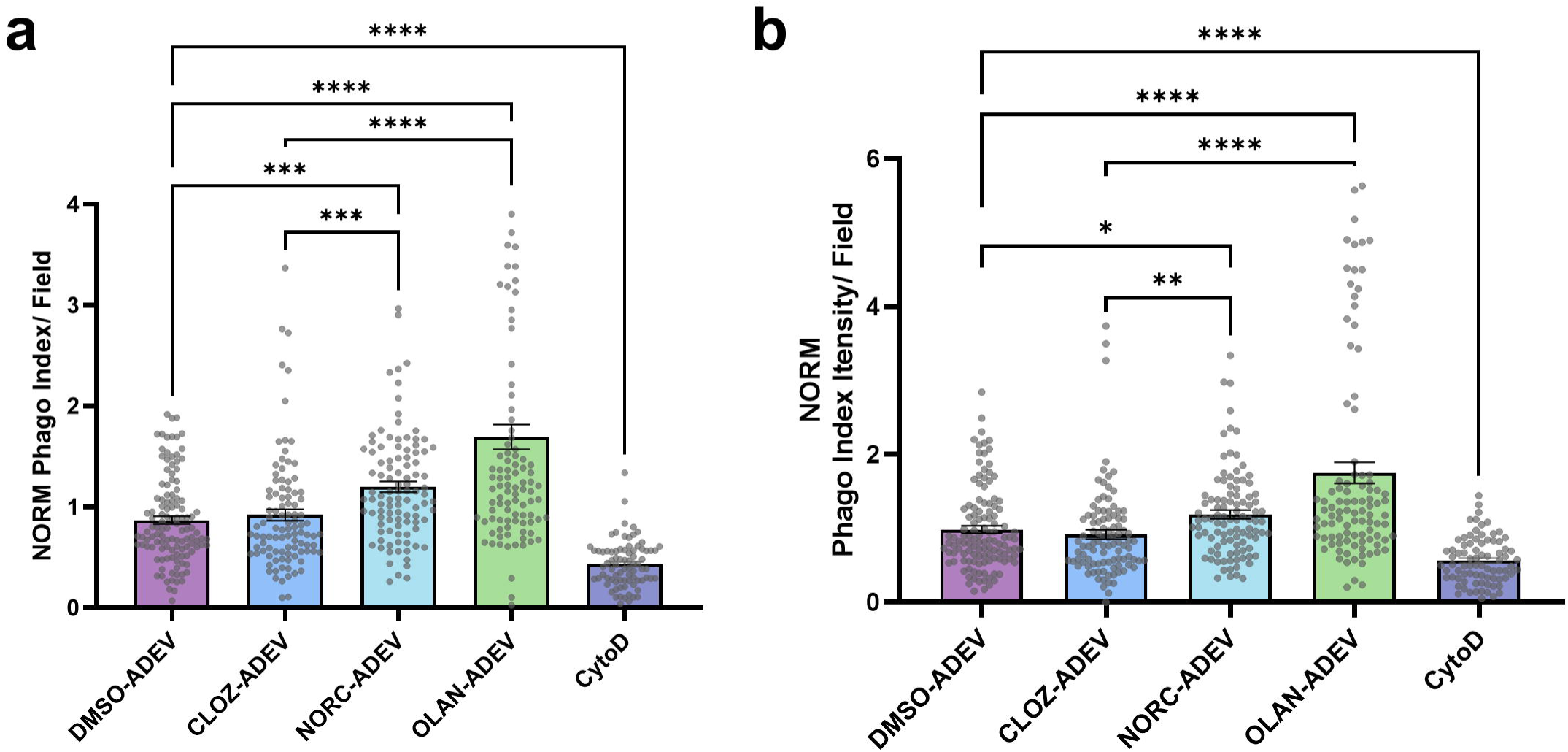
NORC-ADEV and OLAN-ADEV increased synaptosome phagocytosis in piMGLC. **a-b**) ADEVs isolated from atypical antipsychotics norclozapine and olanzapine-treated astrocytes increased phagocytosis index (**a**) and phagocytosis (Phago) index intensity (**b**), with CytoD as control. Kruskal-Wallis with Dunn’s multiple comparison test. (DMSO-ADEV=119, CLOZ-ADEV=99, NORC-ADEV=100, OLAN-ADEV=98, CytoD=90 fields from 2 independent ADEV harvests and experiments). Bars represent mean ± SEM, ∗*p*<.05, ∗∗*p*<.01, ∗∗∗*p*<.001, ∗∗∗∗*p*<.0001.

### Microglial IBA1 expression increased by ADEVs from antipsychotic-treated astrocytes

Previously, we have shown that microglial morphology correlates with phagocytosis capability^34,55^. More amoeboid microglia and increased IBA1 expression are associated with “activated” states that are more likely to phagocytose synaptosomes^55^. In this study, image-based morphological analysis and IBA1 intensity showed that atypical antipsychotics differentially alter microglia morphological parameters through ADEVs. Despite a weaker effect on phagocytosis, NORC-ADEV significantly reduced microglial cell area and cell solidity (Fig. 5A-C), suggesting smaller and less amoeboid morphology. Compared to DMSO-ADEV, antipsychotic-treated ADEVs including CLOZ-ADEV, NORC-ADEV, and OLAN-ADEV all increased microglial IBA1 expression indicating a more “activated” state (Fig. 5D). These discrepancies between phagocytosis and morphology changes suggest ADEVs can alter various microglial biology in distinct ways despite sharing common receptor targets in astrocytes. Lysosomal marker CD68 is upregulated in immune-activated microglia^56^, and often used as a marker for “activated” phagocytic microglia. OLAN-ADEV-treated microglia show elevated intensity of CD68, corresponding to increased phagocytosis (Fig. 4A-B, 5E).

**Figure 5.**
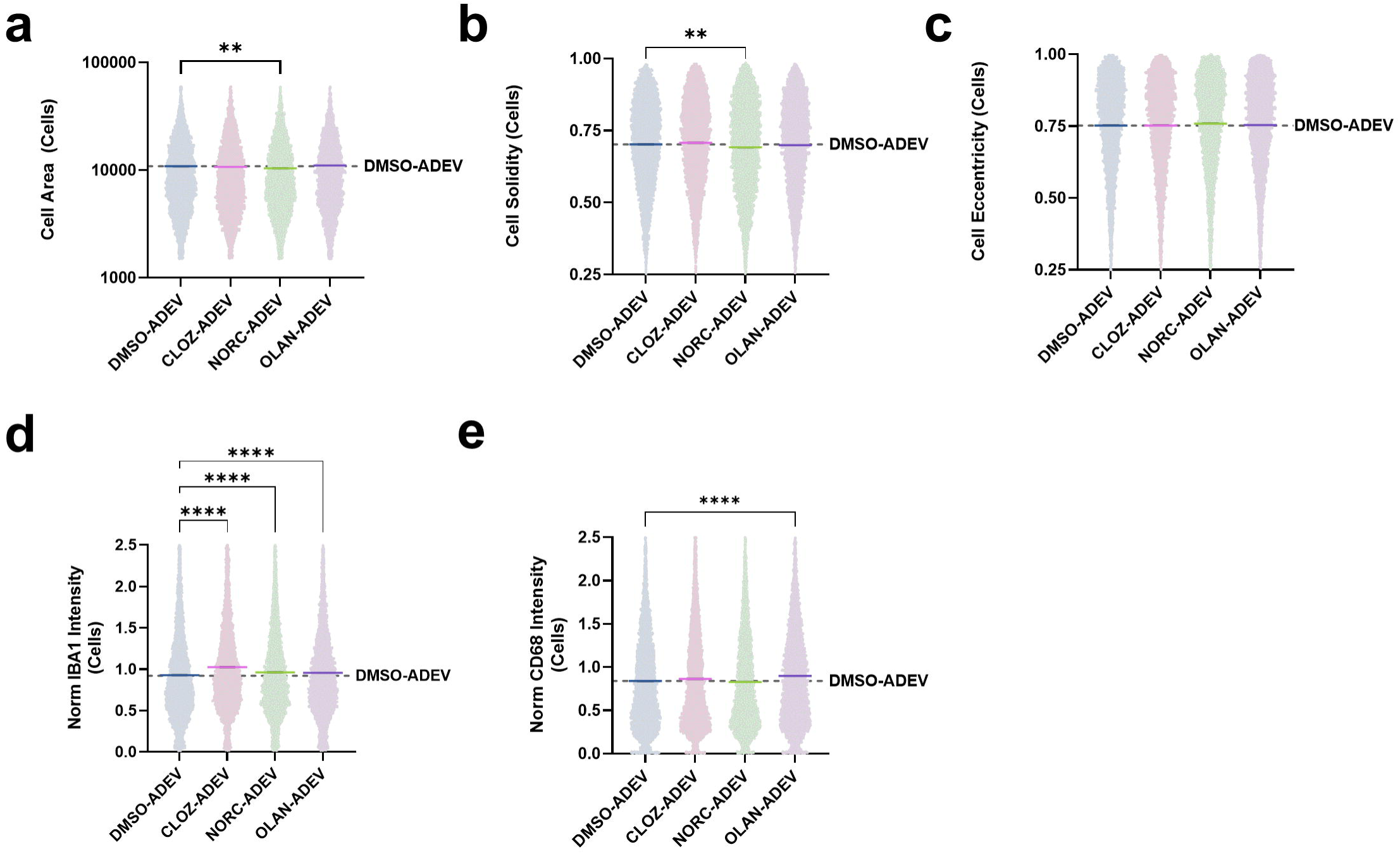
Antipsychotics treated-ADEV have differential effects on microglia morphology and IBA1 expression. **a**) piMGLC size reduced by NORC-ADEVs. **b**) Reduced cell solidity by NORC-ADEVs, and (**c**) no change in cell eccentricity. **a-c**) DMSO-ADEV=7155, CLOZ-ADEV=7519, NORC-ADEV=7524, OLAN-ADEV=6922 cells from 2 independent experiments from at least 6 wells. **d**) increased IBA1 intensity across treatments. DMSO-ADEV=6955, CLOZ-ADEV=7221, NORC-ADEV=7298, OLAN-ADEV=6703 cells from 2 independent experiments. **e**) OLAN-ADEVs increased CD68 expression in piMGLCs. DMSO-ADEV=7097, CLOZ-ADEV=7415, NORC-ADEV=7460, OLAN-ADEV=6888 cells from 2 independent experiments. Outliers removed based on 1% Grubbs’ test. Kruskal-Wallis test with Dunn’s multiple test. Bars represent mean ± SEM, ∗*p*<.05, ∗∗*p*<.01, ∗∗∗*p*<.001, ∗∗∗∗*p*<.0001.

### ADEVs isolated from olanzapine-treated astrocytes reduced microglia motility, whereas clozapine-treated ADEV increased microglia motility

Changes in microglial motility can indicate how microglia survey and physically interact with other cells in the brain parenchyma^42^. In our study, motility assays were used to measure the general surveillant movement of microglia *in vitro*. Using real-time live imaging, we quantified the accumulated distance and velocity of individual microglia-like cells. CLOZ-ADEVs increased microglia velocity and accumulated distance traveled, whereas NORC-ADEVs showed no difference compared to control, and reduced distance traveled compared to ADEVs from clozapine-treated astrocytes (Fig. 6A-B). Increased velocity and accumulated distance traveled indicate microglia are more “hyperactive” and more sensitive to changes in the extracellular environment. Compared to ADEVs from other antipsychotic-treated astrocytes, OLAN-ADEV-treated microglia show the least distance traveled and reduced velocity (Fig. 6B). This suggests different antipsychotics have differential effects on microglia motility through ADEVs.

**Figure 6.**
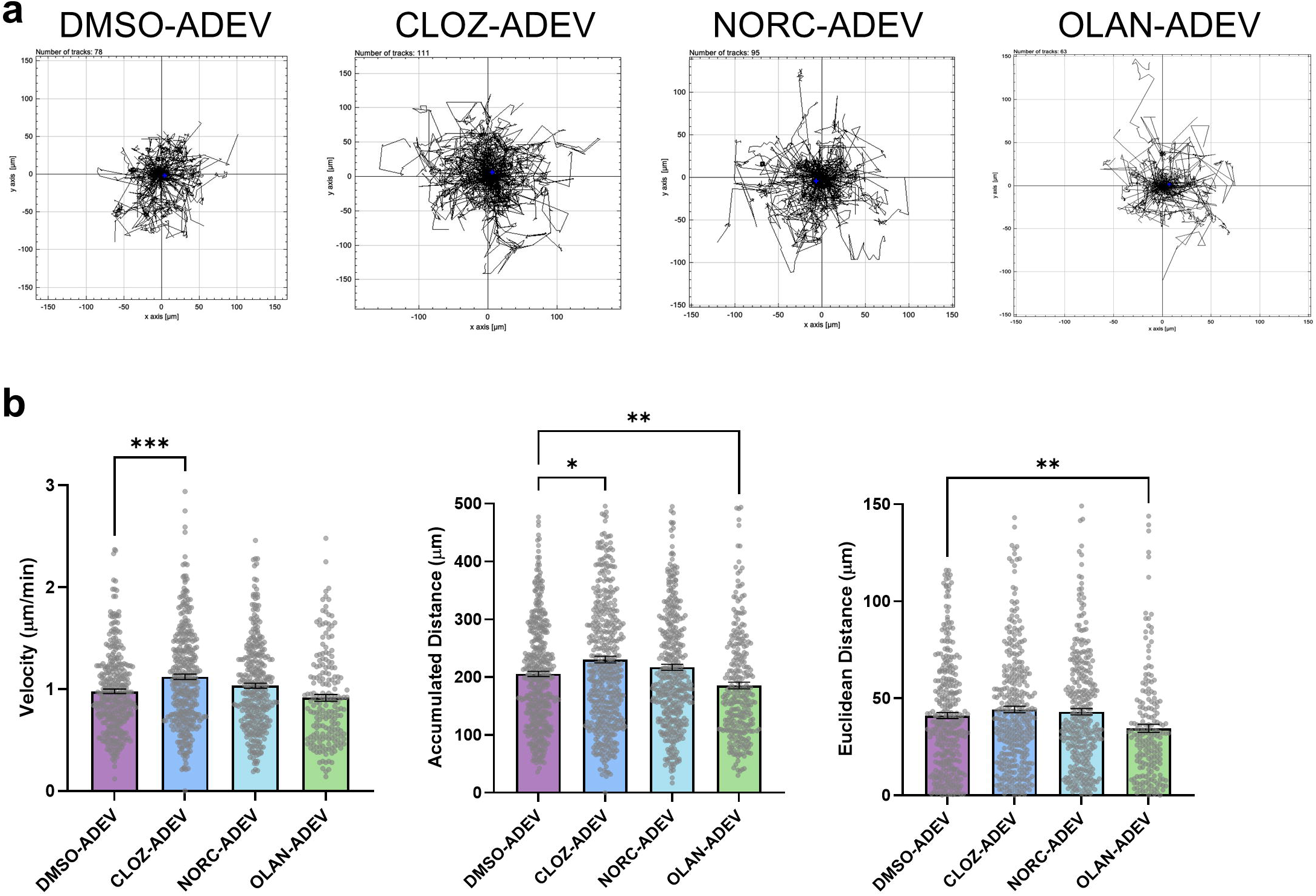
CLOZ-ADEV and OLAN-ADEV have opposing effects on microglial motility. **a**) Cell tracking motility traces of antipsychotic-ADEV treated piMGLCs over a total of 5 hours. **b**) CLOZ-ADEVs and OLAN-ADEVs altered microglia in distinct ways. Kruskal-Wallis with Dunn’s multiple comparison test. (DMSO-ADEV=329, CLOZ-ADEV=346, NORC-ADEV=320, OLAN-ADEV=188 cells from at least 3 independent batches across at least 2 wells per batch). Bars represent mean ± SEM, ∗*p*<.05, ∗∗*p*<.01, ∗∗∗*p*<.001, ∗∗∗∗*p*<.0001.

## Discussion

EVs have emerged as an important communication method between cells in the brain. Astrocytes are one of the most abundant cell types and major producers of EVs in the brain^12,25,57^. Many recent studies highlight the role of astrocytes and ADEVs in contributing to neurodevelopment through EV-mediated glial-to-neuron communication^20,21,46,58^. A previous study also found that EVs secreted from ATP-stimulated microglia increased astrocytic expression of proinflammatory markers including IL-1β, IL-6 and TNFα^15^. We found that, in addition to mediating neuronal functions, ADEVs can also modulate microglial functions and alter their morphology under both homeostatic and inflammatory conditions. The dose-dependent effect of ADEVs on microglial phagocytosis and IBA1 expression highlights that ADEV abundance is required to alter microglial functions.

Previous studies suggest ATP and IL1β stimulants differentially alter neuron physiology based on ADEV cargo content^20,46^. Chaudhuri et al. found ATP-ADEVs increased the frequency of neuronal firing and its ADEV content was enriched with miRNAs associated with neurotrophin signaling pathway and proteins associated with neurite outgrowth, regulation of synaptic transmission pathways; whereas IL1β-ADEVs had the opposite effect of reducing firing frequency and its content was enriched with proteins associated with immune response pathways^20,46^. Our results further suggest that stimulation of astrocytes with IL1β or ATP can differentially affect microglia functions via EV secretion, which is consistent with previous studies that show astrocytes respond to inflammatory conditions by altering miRNA and protein content in ADEVs to promote synaptic stability and control neuronal excitability^20,46,48^. Previous studies have shown that ADEVs contain heat shock protein 70 (HSP70), which is known to induce microglia activation^21,59^. During neurodevelopment, microglia associated with proinflammatory profile selectively uptake neonatal astrocytic EV, suggesting EV uptake is dependent on cell activation state as well^60^. In this study, we did not find that IL1β altered secretion of ADEVs, though we found ATP stimulation reduced ADEV secretion, suggesting different activation states of astrocytes could alter EV secretion and ATP-ADEVs led to more ADEV load index per PBMC-microglia cell than other treatments.

Antipsychotics share similar chemical structures but can impact cell functions very differently due to different biological targets – differences that may be apparent in their heterogeneous adverse effect profiles^61–64^. Despite no apparent effects on astrocytic markers, antipsychotics altered microglial marker expression and cellular morphology via ADEVs, suggesting these drugs may have an indirect impact on glial cells via EVs. In this study, we found that ADEVs from antipsychotic-treated astrocytes increased microglial phagocytosis and IBA1 expression compared to DMSO-ADEV treatments on PBMC-derived microglia-like cells. CLOZ-ADEVs increased microglial motility and IBA1 expression, but had no effect on phagocytosis, suggesting that clozapine alters microglial surveillance within the environment and relative activation state. On the other hand, NORC-ADEVs had no impact on motility, but increased phagocytosis and reduced cell size and solidity, indicating a different active state that targets nearby cells and environment. OLAN-ADEVs had the most significant increase in microglial phagocytosis and CD68 expression but had a very subtle impact on motility, suggesting that olanzapine pushed microglia toward a more targeted phagocytotic state. Koskuvi et al. showed clozapine did not significantly alter microglial phagocytosis^65^, which is consistent with lack of CLOZ-ADEV effect on PBMC-derived microglial-like cells in this study. The differential effects on cellular morphology, lysosomal CD68 marker, and motility due to different antipsychotic-treated ADEV conditions highlight the need to further investigate antipsychotics to evaluate and compare how they differ in mechanisms on the cellular level.

Additional studies may be required to examine other labeled substrates to examine whether antipsychotics affect the identification of specific substrates or general phagocytosis capabilities. Future studies could examine the EV content of antipsychotic-treated EVs including nucleic acids and protein cargo. Recent studies show antipsychotics can alter the miRNA content of EVs and regulate synaptic biology in human cortical neurons ^27,66^. Psychosis-altered and glial-enriched miRNA can mediate neuronal NMDA receptor gene expression via exosomes, which are small secretory microvesicles between 30-150 nm mainly produced from the multivesicular bodies of the endosomal pathway, and be reduced by antipsychotics^27^. Two-day treatment with olanzapine and haloperidol reduced the expression of miR-223 in murine neurons and astrocytes, and increased miR-223 astrocytic exosomal secretion, suggesting antipsychotics can affect EVs based on cell type^27^. Furthermore, Funahashi et al. found differentially expressed exosomal miRNA derived from plasma exosomes of treatment-resistant schizophrenia patients compared to treatment-responsive patients^67^. Because of their ability to cross the blood-brain barrier, EVs may find application both as diagnostics and therapeutic agents for psychiatric disorders. Engineered EVs and EV-mediated drug delivery for targeted treatments in specific cells could be developed by genetically or chemically changing EV membrane molecules and cargo content^26^. In a similar fashion, understanding how olanzapine and clozapine can improve microglial function through EVs and modifying microglial miRNA content via EV delivery can potentially provide an alternative therapeutic approach for treatment-resistant schizophrenia in a more cell-targeted manner.

## Conclusion

To directly quantify various cellular functions, we utilized primary cellular models for both astrocytes and microglia cell models. A strength of this design is that we were able to isolate pure ADEVs and examine their direct impact on microglial-like cells using monoculture models separately. We were able to investigate microglia functions across multiple parameters under different ADEV conditions and identify key differences between different ADEVs isolated from antipsychotic-treated astrocytes that support our hypothesis that antipsychotics can alter microglia function via ADEVs. Future studies can examine how schizophrenia risk genes can contribute to antipsychotics-treated ADEV response in patient-derived cells, and further investigate using *in vivo* animal models or complex organoid cellular models to study glia-to-glia EV communication under different stimulants relevant to schizophrenia or other psychiatric disorders. This study is a proof-of-concept to study how drugs can alter microglia functions via ADEVs as an indirect communication method.

## Resource Availability

Additional data that support the findings of this study will be available from the corresponding authors upon reasonable request. Additional assay details are available in Supplemental Material and Methods.

## Acknowledgements

We would like to acknowledge training and access to Nanosight LM10 at Massachusetts General Hospital Exosome Quantification Research Core.

## Author Contribution

HY and RHP conceptualized, designed, and developed this study. HY, YS, LTM, and JJB performed experiments. HY and YS analyzed the data. RHP and SDS supervised the research and consulted with data interpretation. HY, SDS and RHP wrote the manuscript. All authors have read and approved the manuscript for publication.

