## Supplemental Data for "Atypical antipsychotics alter microglial functions via astrocyte-derived extracellular vesicles"

| Antibody | Species | Vendor (cat#) | Dilution |
|---|---|---|---|
| **IBA1** | Chicken monoclonal | Synaptic Systems (234009) | 1:1000 |
| **S100b** | Mouse monoclonal | BioLegend (676604) | 1:100 |
| **ALDH1L1** | Rabbit polyclonal | Abcam (ab190298) | 1:100 |
| **GFAP** | Chicken polyclonal | Novus Biology (NBP1-05198) | 1:1000 |
| **Chicken secondary** | Goat-Alexafluor647 | Invitrogen (A21449) | 1:1000 |
| **Mouse secondary** | Donkey-Alexafluor488 | Invitrogen (A21202) | 1:1000 |
| **Rabbit secondary** | Donkey-Alexafluor555 | Invitrogen (A31572) | 1:1000 |

**Supplemental Table 1.** Primary and secondary antibodies used for immunocytochemistry and confocal microscopy imaging

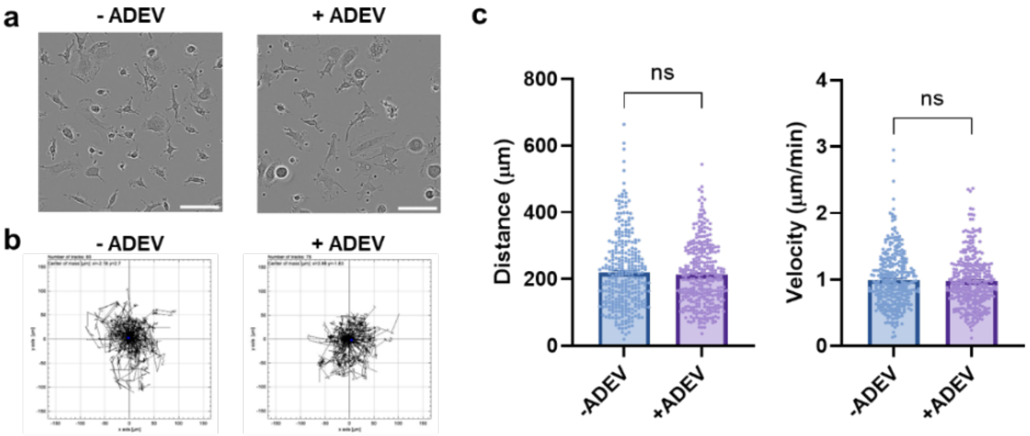


**Supplemental Figure 1.** ADEV does not alter microglial motility. **a**) Representative images for piMGLCs. **b**) piMGLC motility track traces treated with (+ADEV) and without (-ADEV) astrocyte-derived EVs. **c**) No significant changes in total accumulated distance travelled (left) and velocity (right) between -ADEV and +ADEV. Each dot represents individual cells. (Mann-Whiteny test *n*=316-329 cells from at least 3 independent batches across at least 2 wells per batch).

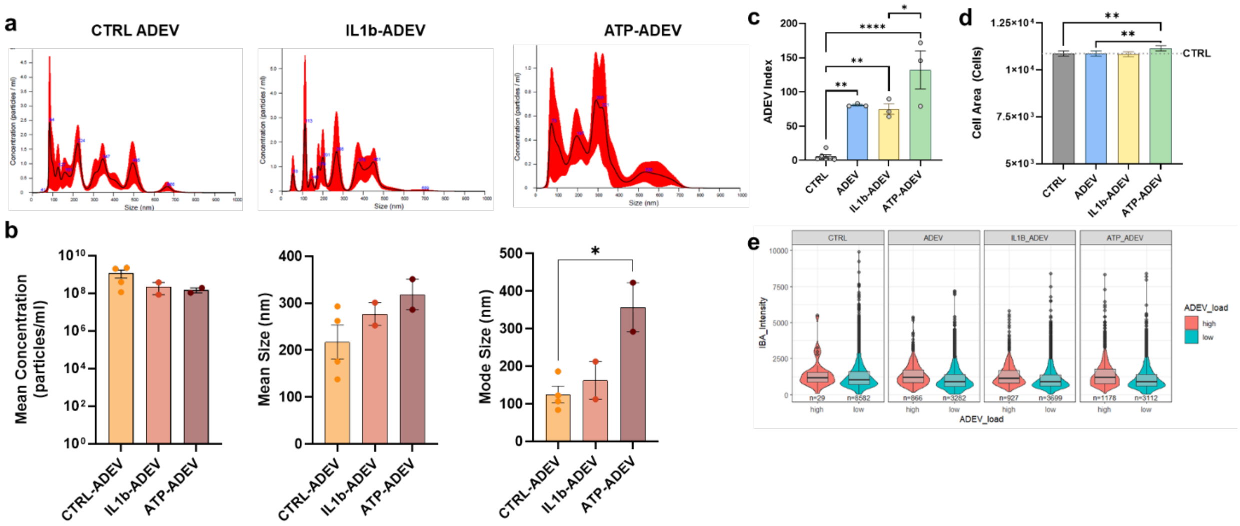


**Supplemental Figure 2.** IL1β and ATP-ADEV characterization show ATP reduced mean ADEV concentration and increased piMG uptake. **a**) Representative NTA traces of diluted ADEVs. **b**) Mean concentration (left), size (middle), and mode (right) of ADEVs across different conditions including IL1β and ATP stimulation. Tukey's multiple comparison test (*n*=2-4 ADEV replicates from two independent ADEV isolation experiments). **c**) ADEV index based on area of PKH67 labeled ADEVs per piMGLCs across conditions. Tukey's multiple comparison test (CTRL=6 wells, ADEV, IL1β-ADEV, ATP-ADEV=3 wells). **d**) ATP-ADEV increased cell area. Kruskal-Wallis test with Dunn's multiple comparison test. (*n*=CTRL 4300, ADEV 4148, IL1β -ADEV 4300, ATP-ADEV 4290 cells) **e**) High or low ADEV load based on ADEV index and relative IBA1 intensity. Bars represent mean ± SEM, $*p<.05$, $**p<.01$, $***p<.001$, $****p<.0001$.

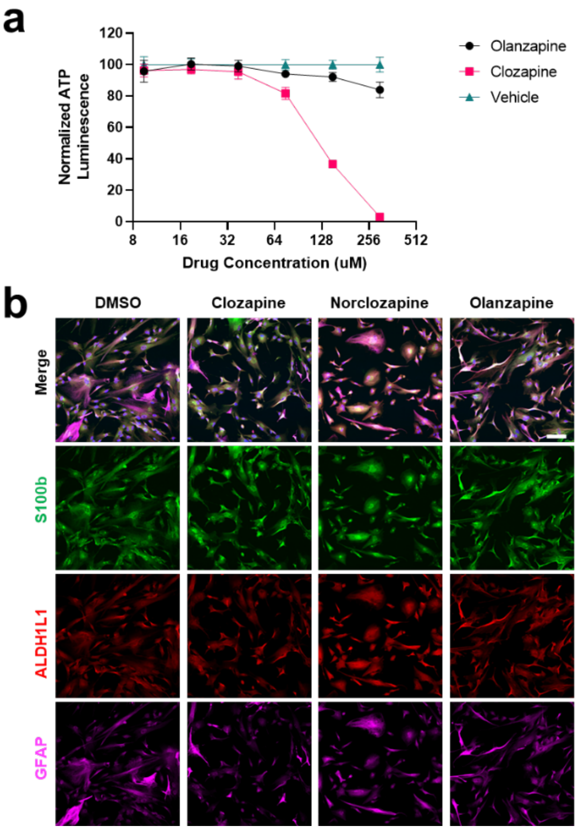


**Supplemental Figure 3.** Clozapine and olanzapine retain astrocytic marker expression. **a**) Normalized ATP luminescence to vehicle (DMSO) using Cell Titre Glo (Lonza) from 9.4 μM to 300 μM of clozapine or olanzapine. *n*=3 wells per treatment. **b**) Representative images of human primary astrocytes (ScienCell HA1800) at P4 with astrocytic markers including S100b, ALDH1L1, and GFAP. Scale bar=100 μm.

### Supplemental methods

*Large scale generation of synaptosomes from iPSC-neural cultures*

Neural cultures were differentiated from iPSC-derived neuronal cultures in T1000 multilayer flasks as previously described[1–3]. Cells were harvested in ice-cold 1X sucrose gradient buffer consisting of 0.32M sucrose, 600mg/L Tris, 1mM NaH3CO3, 1mM EDTA, pH 7.4, HALT protease inhibitor (ThermoFisher, #78442) and homogenized with a Dounce homogenizer. The homogenate was centrifuged at 700 g for 10 minutes at 4˚C. The pellet was resuspended in 1X sucrose gradient buffer and centrifuged at 15,000 g for 15 minutes at 4˚C, then resuspended in 1X gradient buffer and layered on a sucrose gradient (1.2M bottom, 0.85M middle). This mixture was centrifuged at 80,000 g for 2 hours at 4˚C (brake slow). The synaptosome band was collected, centrifuged at 20,000 g for 20 minutes at 4˚C, and resuspended in 1X gradient buffer with 1 mg/mL bovine serum albumin (BSA), protease, and phosphatase inhibitors. Protein concentration was measured using a bicinchoninic acid assay (ThermoFisher, BCA Protein Assay Kit# 23225). Expression of synaptosome markers postsynaptic density 95 (PSD95) and synaptophysin (SYN) was confirmed by western blot previously shown and described[3,4].

*Cell morphology analysis*

All image-based quantification, including phagocytosis, was performed using CellProfiler (Version 4.2.4) and ImageJ on random image fields for unbiased analysis. CellProfiler was used to identify iMGs and quantify cell morphology, and synaptosome phagocytosis. Background subtraction was applied using a 300-pixel Gaussian smoothing kernel. Nuclei were segmented as primary objects in Hoescht images using global Otsu two-class thresholding, with cells identified as secondary objects using the IBA1 stain and the

Watershed-Gradient method. The cytosolic compartment was defined as the cell region excluding the nucleus. Synaptosomes were segmented as primary objects using adaptive Otsu two-class thresholding and masked with the cytosol mask. Background noise was excluded by running the pipeline with control images lacking synaptosomes or treatment with CytoD. Raw data were processed in R and imported into GraphPad Prism for statistical testing.
